# Application of Global Transcriptome Data in Gene Ontology Classification and Construction of a Gene Ontology Interaction Network

**DOI:** 10.1101/004911

**Authors:** Mario Fruzangohar, Esmaeil Ebrahimie, David L. Adelson

## Abstract

Gene Ontology (GO) classification of statistically significant over/under expressed genes is a commonly used to interpret transcriptomics data in functional genomic analysis. In this approach, all significant genes contribute equally to the final GO classification regardless of their actual expression levels. However, the original level of gene expression can significantly affect protein production and consequently GO term enrichment, and genes with low expression levels can participate in the final GO enrichment through cumulative effects. In addition, GO terms have regulatory relationships that allow the construction of a regulatory network that incorporates gene expression levels to better study biological mechanisms. In this report, we have used gene expression levels in bacteria to determine GO term enrichments. This approach provided the opportunity to enrich GO terms across the entire transcriptome (instead of a subset of differentially expressed genes). In the second part, we show a dynamically developed enriched interaction network between Biological Process GO terms for any gene samples. This type of network presents regulatory relationships between GO terms and their genes. We then demonstrate the efficiency of these methods using public data from two important bacterial pathogens as models. We also explain how these methods help us understand potential pathogenesis mechanisms employed by these bacteria.

## Background

The functional genomic changes in bacterial pathogens during disease progression or in emerging highly pathogenic strains are poorly understood. Classifying genes into distinct functional groups through Gene Ontology (GO) is a commonly used and powerful tool for understanding functional genomics and the underlying molecular pathways. However, GO protein enrichment is related to the amount and number of proteins described in that GO, and in eukaryotes mRNA levels are often poorly correlated with protein expression. Bacteria are attractive organisms for GO analysis since they have less Post-transcriptional gene silencing compared to animals and plants (1) with gene expression levels moderately correlated with protein levels (2).

Because of the lack of specific resources for GO analysis in bacteria, we recently developed Comparative GO, a PHP based web application for statistical comparative GO and GO-based gene selection in bacteria (3). Comparative GO has the potential to provide a comprehensive view of bacterial functional genomics by categorizing genes into a limited number of annotated GO groups (3,4).

Another major advantage of GO analysis is developing quality-based gene selection strategies compared to the common approach of gene selection in bacteria which is solely based on the level of gene expression (quantity based gene selection) (3,4). It should be noted that expression level alone cannot be used as a sole index of gene significance because some genes with lower expression levels (such as transcription factors) play a prominent role in bacterial systems biology (3,4). An integrative approach, combining quality-based metrics such as GO classification, promoter analysis, and network construction in conjunction with quantity-based gene selection criteria provides a more robust approach for identifying key bacterial genes and describing bacterial systems biology. Such an approach can contribute to the discovery of genes associated with specific function(s) for investigation as novel vaccine candidates or pathways for pharmacological targeting.

Biological process GO terms are analogous to genes because they have regulatory relationships with each other that can be used to construct a directed acyclic network. Compared to common gene networks, GO regulatory networks can identify key functional genomics based interactions in a broader sense. Classifying a large number of genes in a small number of GO classes and visualising the GO networks can significantly decrease the network complexity and, more importantly, offers a new approach for gene selection by considering the genes which contribute to central nodes in GO networks. To our knowledge there is no tool and methodology currently available to dynamically construct GO regulatory networks.

The common approach in transcriptome experiments is that GO analysis is carried out on a short list of genes with statistically significant differential expression (up/down regulated) (5–7). In this approach, all selected genes contribute equally in the final GO classification regardless of their actual expression levels.

The major drawback to this approach is that the original levels of gene expression can significantly affect protein production and consequently actual GO term enrichment. In addition, even genes with low or statistically non-significant expression levels can participate in final GO enrichment through cumulative effects.

In this report we show for the first time how gene expression levels in bacteria can be used to determine GO term enrichments. By using gene expression levels as coefficients, we also took into account the impact of non-significantly expressed genes in GO enrichment. This approach provided the opportunity to enrich GO terms from the entire transcriptome genome (instead of samples of a short list of genes) and enabled us to compare GO terms of transcriptomes across multiple biological conditions. In order to achieve this, we enhanced our recently developed web server, Comparative GO (3) available at http://genomes.ersa.edu.au/BacteriaGO/. To enable analysis of very large gene sets such as from a whole genome, we implemented cache technology to improve web server performance. We also integrated robust non-parametric chi-square based tests into our web application to enable genome scale GO based comparison of gene expression.

We applied our new methods to two important bacterial pathogens, *streptococcus pneumonia* and *Salmonella Enteritidis* in order to unravel the global, transcriptome based, GO pattern of *streptococcus pneumonia* during infection of host tissues and breaching of tissue barriers as well as the comparison of low and highly pathogenic *Salmonella Enteritidis* strains (8).

In the second part of this study we describe the implementation of GO based gene selection and GO network discovery. We show for the first time a dynamically constructed and enriched interaction network between Biological Process GO terms for any given bacterial gene sample. To this end, GO relationships were extracted from the Gene Ontology database (9,10), and used to build a directed acyclic graph (DAG). To visualise the final DAG, we used the Cytoscape web browser plug-in (11). We used our *streptococcus pneumonia* and *Salmonella Enteritidis* data sets as case studies for this method.

## Material and Methods

### Incorporation of gene expression levels into GO analysis

The system accepts any type of expression level such as microarray fold-change data and RPKM counts of RNA-Seq data. In all cases, for each gene, one coefficient is estimated based on its expression level within the sample or within the genome. If we want to perform comparative GO analysis on n sample of genes, and the expression level of gene i in list j is e_ij_ and also given that the smallest expression level across n samples is denoted by e_min_, then the coefficient of gene i in list j (C_ij_) is estimated as:

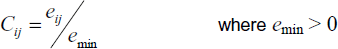

Furthermore, if GO term t in sample r is associated with genes G1r…Gmr, then the protein enrichment level of GO term t in sample r (PEtr) is estimated as:

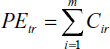

We have implemented these methods on our server with PHP.

### Hypothesis Testing Tool

We implemented a tool to test the hypothesis of a significant difference between 2 genome/sample GO term distributions. Specifically, we implemented a Chi-Square test for 2 samples and we compared it with the Kolmogorov–Smirnov test using the R-statistical package(12). These two methods are both non-parametric and are suitable for comparing 2 lists of paired numbers like GO term enrichment values for 2 samples.

### GO regulatory Network Construction

Regulatory relationships (up/down regulation) were extracted between Biological Process terms from the Gene Ontology database available at http://www.geneontology.org/ (9,10). We stored these relationships in our internal database (3). For any given gene sample, our application builds a GO DAG (Directed Acyclic Graph) network, based on regulatory relationships.

In order to infer new relationships from explicit relationships we expanded the initial GO network to include parental nodes; then, new relationships were inferred from relationships between parental GO nodes to the nodes in the network. Figure 1 depicts a simple GO regulatory network, where grey nodes represent the GO terms related to the sample, and the relationships between GO terms are depicted by green arrows. As we can see at the top of the graph, there is a relationship between parental GO terms 2 and 3. Accordingly, we inferred 3 new relationships between nodes 4, 5, 6 and node 7, depicted as green dotted arrows. The final enriched network can describe novel regulatory relationships between GO terms and consequently between their associated genes.

**Figure 1.**
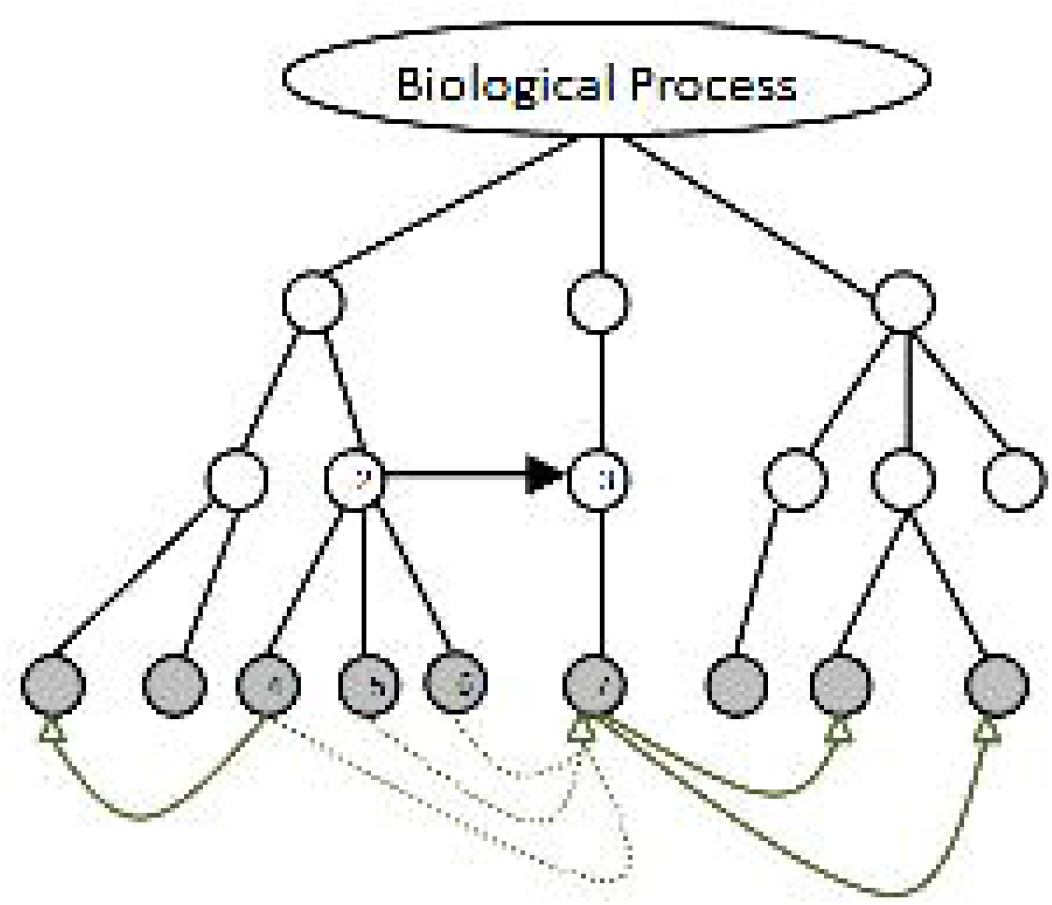
Schematic presentation of a simple GO regulatory network. Grey nodes represent GO terms related to the sample and the relationships between GO terms are depicted as green arrows. Parental GO nodes 2 and 3 have a relationship that can be extended to children GO nodes 4, 5, 6 and node 7, depicted as green dotted arrows.

### Web Application Enhancements

Methods and algorithms were implemented in our web application (3) using PHP 5 and PostgreSQL. Because of the additional functionality to analyse the GO distribution of all expressed genes within a genome (global transcriptomics), significant memory and processing resources were required by the Apache web server. To enhance performance and husband system resources we implemented file based caching technology to cache the whole genome GO graphs. When a GO graph was built for the first time, subsequent references to that GO graph, even by other users, were instantaneous. For a better user experience in web application pages where long running tasks were performed, we used Ajax technology to implement task progress bars.

### Visualising the GO interaction network

In order to visualize the enriched GO regulatory network, we used the Cytoscape (11) flash player plug-in for web. We initialized and used this component through JavaScript. Cytoscape contains advanced dynamic network customization options such as zooming, network filtering, node re-locating, node and edge re-sizing, and colour scheming. These user-friendly options allow developers and users to dynamically change the look and feel of the network.

### Case study data

To demonstrate the biological application of our new methods in global transcriptome GO analysis and GO network analysis, data from 2 previous gene expression experiments were used. *Streptococcus pneumonia* and *Salmonella enteritidis* were selected since they are responsible for high morbidity, mortality, and infection worldwide and have been well studied.

The first data set (4) was two colour microarray data from *Streptococcus pneumonia* in vivo derived RNA samples, where the relative expression of each gene in one niche was calculated in comparison to expression in the previous niche when bacteria moved from nose → lungs → blood → brain. The relative expression of all 2236 genes of *Streptococcus pneumonia* during the course of infection are presented in Additional File 1 (lung versus nose), Additional File 2 (blood versus lung), and Additional File 3 (brain versus blood). Additional files are in MS Excel worksheet format.

The second case study (8) was RNA-Seq global transcriptome data from 6 strains of *Salmonella enteritidis*, where 3 highly pathogenic strains and 3 low pathogenic strains were compared. The average whole genome expression of (4402) genes of the 3 highly pathogenic strains is presented in Additional File 4. While Additional File 5 contains the average expression of the 3 low pathogenic strains. The goal of this analysis was to identify significantly different GO terms between highly and low pathogenic strains of bacteria using *Salmonella enteritidis* as a model.

For GO network analysis, in case study 1, the 30 highest over expressed genes in *Streptococcus pneumonia* during infection in lung versus, blood versus lung and brain versus blood were used (Additional File 6). Also, in case study 2, 18 genes with the highest fold change in expression levels between highly pathogenic strains versus low pathogenic strains are presented in Additional File 6.

## Results

### Introduction of gene expression levels into GO analysis

Addition of expression level data with GO term data provided the opportunity of (a) quantifying more accurate GO enrichments, (b) extending analysis coverage from sample-wide to genome-wide, and (c) developing statistical tests for comparison of GO distributions across transcriptomes. Considering the influences of all expressed genes in functional genomics, even those with low levels of expression, could possibly increase the accuracy of the analysis in prokaryotes.

### GO regulatory network

GO regulatory networks for a sample of genes initially present three types of information: regulatory relationships between GO terms and their associated genes depicted by directed edges of the graph, enrichment levels of GO terms that are proportional to nodes’ diameter of graph and finally, the genes associated with each GO term.

Furthermore, network topology revealed GO groups and their genes that had the highest number of interactions with other groups. Specifically, genes located at the centre of the network were selected as good candidates for further experiments and gene discovery. In addition, the enrichment levels of GO terms that were proportional to the size of the nodes in the graph were in accordance with the regulatory relationships between GO terms.

### Case studies

As case studies, we used publicly available two colour microarray and global transcriptomics data of two important bacterial pathogens, *Streptococcus pneumonia* and *Salmonella enteritidis* respectively. For each bacterium, 2 types of analysis were carried out: transcriptome based GO enrichment and GO network discovery. In Streptococcus pneumonia, all expressed genes were subjected to GO analysis in order to characterise functional changes in *Streptococcus pneumonia* during the course of infection. Then, using a selection of significantly up-regulated genes during infection in each tissue, GO networks were constructed to identify the central GO node and the key genes associated with the central GO node. In the *Salmonella enteritidis* case study, we first compared transcriptome GO enrichment levels between highly pathogenic and low pathogenic strains to highlight GO functional groups correlated with pathogenicity. We then constructed the GO network using the genes that were significantly more highly expressed in pathogenic strains.

#### Case Study 1: Changes in the transcriptome GO during Streptococcus pneumonia from nose → lungs → blood → brain

After downloading microarray data (4) from the NCBI GEO database for *Streptococcus pneumonia*, we selected data of strain WCH43 after 72 hours infection across 4 different tissues. We estimated the geometric means of the fold-change for each gene in the genome. The result was 3 genome-wide lists (Nose vs. Lung, Lung vs. Blood and Blood vs. Brain) each containing 2236 genes along with their mean fold-changes (Additional File 1, 2 and 3). These lists were submitted to the web server.

First, we used the pie chart visualisation to determine GO term proportions (protein enrichment distribution percentage) at different levels of the GO tree. GO term proportions of some GO groups didn’t change across multiple tissues. Hence, the GO term proportions of 3 genome-wide lists were mutually compared by Kolmogorov–Smirnov test and the calculated p-values are presented in Table 1.

**Table 1.** Comparison of genome-wide GO enrichment levels by Kolmogorov–Smirnov test during the infection course of *Streptococcus pneumonia* from nose → lungs → blood → brain

|  | **Biological Process** | **Molecular Function** | **Cellular Components** |
| --- | --- | --- | --- |
| **(Lung vs. Nose ~ Blood vs. Lung)** | P=0.01 | P=0.01 | Not significant |
| **(Blood vs. Lung ~ Brain vs. Blood)** | Not Significant | P=0.01 | Not significant |

Table 1 suggests that Cellular Components GO enrichment proportions did not change at all during the course of infection. Interestingly, when bacteria moved from blood to their final destination (brain), the overall proportions of Biological Process GO terms did not change.

We then produced a tabular report of the last level (most detailed) of the GO tree. From a large list of GO terms, this report highlighted GO terms that were consistently up/down regulated. Surprisingly, in this study only identified a few such GO terms (Figure 2). GO terms with upward or downward arrows had consistent up/down expression patterns. The continuously up regulated GOs were “barrier septum assembly” and tryptophan synthase activity which are involved in propagation of *Streptococcus pneumonia*. This result confirmed a known, experimentally verified mechanism in this organism (4). The list of genes in each GO is also presented to assist with GO based gene selection. GOs such as “histidine biosynthesis process” and “amidase activity” were down regulated. This report also highlights GO terms with more than 4 fold average fold-change.

**Figure 2.**
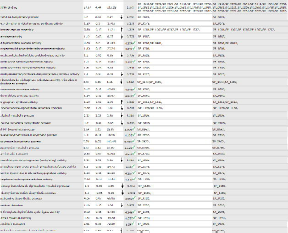
Amended “Table report” which lists consistently up and down regulated GO terms and also GO terms with more than 4 times change in protein enrichment.

#### The GO regulatory network during Streptococcus pneumonia infection from nose → lungs → blood → brain

The GO network during movement of *Streptococcus pneumonia* from nose to lung is presented in Figure 3A. Upon inspection, regulation of transcription (Gene Ontology ID: 6355) is a central node in the network and SP_0798 is the only component of this GO network. Interestingly, the GO group (regulation of transcription) governed by SP_0798 plays a key role in breaching the brain-blood barrier and infection of brain tissue. We previously demonstrated that the SP_0798 transcription factor positively regulates the Sp-0927 transcription factor and activates a sub network through interaction with proteins such as SP_0797, SP_0084, SP_2083, SP_1226, and SP_0799 (4). The SP_0798 sub network is one of the key sub networks conferring high virulence to *Streptococcus pneumonia* (4).

**Figure 3.**
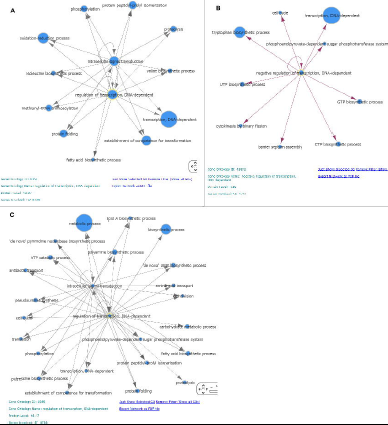
GO regulatory network constructed based on differentially expressed *Streptococcus pneumonia* genes in (A) Lung versus Nose (B) Blood versus Lung (C) Brain versus Blood.

When comparing lung-nose niche expression patterns, the SP_0798 governed GO has interactions with GOs such as: “phosphorylation”, “fatty acid biosynthesis process”, “establishment of competence for transformation” and “oxidation-reduction process”. The “establishment of competence for transformation” GO (SP_0798 gene) can play a significant role in the translocation of *Streptococcus pneumonia* from nose to lung.

Figure 3C showed that the SP_0798 governed GO (Gene Ontology ID: 6355) had a considerable number of regulatory effects in the brain-blood comparison. The brain is the final destination of *Streptococcus pneumonia* WCH43 where it causes meningitis. SP_0798 activated different GO groups such as “metabolic process”, “establishment of competence for transformation”, “phosphorylation” and “antibiotic transport” while reaching and infecting the brain. Activation of “antibiotic transport process” helps *Streptococcus pneumonia* resist antibiotics.

It was previously (4) known that in meningitis-inducing strains of *Streptococcus pneumonia* such as WCH43, relative global gene expression significantly decreased in blood compared to the previous niche (lung) or the subsequent niche (brain). Interestingly, the GO network shown in Figure 3B helps illustrate the underlying mechanism of this global down regulation and shows that Gene Ontology ID 45892 (“negative regulation of transcription, DNA-dependent”) governed by SP_1713 transcriptional repressor NrdR is central to this relative decrease in expression. Gene Ontology ID 45892 has interactions with “CTP/GTP biosynthesis process”, “barrier septum assembly” (involved in propagation), “cytokinesis binary fission”, and “tryptophan biosynthesis process” (Figure 3B). The SP_1664 protein is involved in barrier septum assembly. SP_1813, SP_1814 and SP_1815 proteins participate in tryptophan biosynthesis process.

Discovery of the Gene Ontology ID 45892 (“negative regulation of transcription, DNA-dependent”) governed by SP_1713 and its considerable influence in suppression of genes opens a new avenue for the investigation of treatment for blood stream-based diseases such as Bacteremia and Sepsis.

#### Case Study 2: Comparison of whole transcriptome based GO enrichment between low and highly pathogenic *Salmonella enteritidis*

We collected RNA-Seq data for 6 strains of low and high pathogenic *Salmonella enteritidis* (8) including 3 low pathogenic strains and 3 highly pathogenic ones. We averaged the RPKM counts for each gene of the 3 low pathogenic strains and created a single list of genome expression levels. We did the same for the 3 highly pathogenic strains (Additional File 4 and 5). After submission of both gene lists (4402 genes for each one) to the web server, we used the pie chart to visualise the GO term proportions and navigate the GO term tree. The comparison revealed very similar GO proportions at nearly all levels of the GO tree. This encouraged us to perform hypothesis tests to compare the GO enrichment proportions between low and highly pathogenic strains. Table 2 shows the result of this comparison for Biological Process, Molecular Function, and Cellular Components.

**Table 2.**
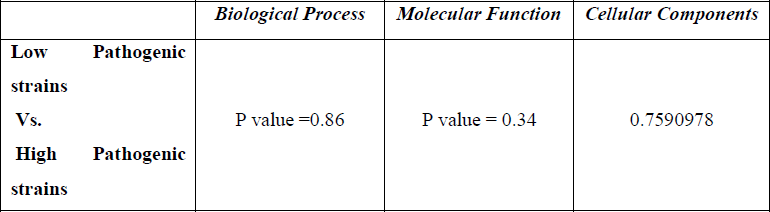
Comparison of genome wide GO enrichment levels of low pathogenic strains of *Salmonella enteritidis* versus high pathogenic strains by Kolmogorov–Smirnov test

|  | **Biological Process** | **Molecular Function** | **Cellular Components** |
| --- | --- | --- | --- |
| **Low Pathogenic strains Vs. High Pathogenic strains** | P value =0.86 | P value = 0.34 | 0.7590978 |

Based on a 0.05 level of significance for our tests, Table 2 indicates that there is probably no significant difference in GO protein enrichment proportions between low and highly pathogenic strains of *Salmonella enteritidis* bacteria (null hypothesis accepted). This is a significant finding that demonstrates that the change from low pathogenic strain to highly pathogenic strain is not associated with a global shift in GO terms. However, as seen below, a shift in a subset of GO terms can be associated with higher pathogenicity.

#### GO regulatory network changes between high and low pathogenic strains of Salmonella enteritidis

A list of the most differentially expressed genes - with greater than 10 fold change - were submitted to the Web server (Additional File 6), including fljB, SEN1084, motA, flgK, cheA, invF, invA, invG, fliD, prgH, osmY, ipB, sipC, yeaG, sipA, dps, yjbJ, and bfr. The resulting GO network is presented in Figure 4.

**Figure 4.**
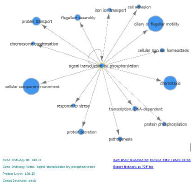
GO regulatory network based on 18 genes with significant differential expression levels in highly pathogenic versus low pathogenic *Salmonella enteritidis* strains.

Interestingly, the GO term “signal transduction by phosphorylation” (Gene Ontology ID: 23014) is central in the overrepresented GO expression network of highly pathogenic *Salmonella enteritidis* strains. The protein kinase encoded by cheA is the sole component of “signal transduction by phosphorylation process”. This shows that higher pathogenicity in *Salmonella enteritidis* appears to be associated with increased signal transduction and phosphorylation. We speculate that up regulating GO “Signal transduction by phosphorylation” may allow *Salmonella enteritidis* to more rapidly sense environmental changes and activate more genes through stronger phosphorylation activity. “Response to stress”, “iron ion transport” (bfr gene), “pathogenesis”, “transcription DNA dependent”, “protein phosphorylation” (yeaG gene) and “chemotaxis” are the other GO terms which are differentially expressed in the highly pathogenic strain.

### Commonality between GO Regulatory Networks of Case Studies

Selection of *Streptococcus pneumonia* during the course of infection in nose, blood, and brain of host allowed us to apply whole genome based GO enrichment and GO in a study of tissue-based pathogenesis and breaching of host barriers by a pathogen. In addition, comparative study of GO enrichment and GO network between highly pathogenic and low pathogenic strains of *Salmonella* allowed us to investigate mechanisms involved in the generation of highly pathogenic strains using GO networks.

Go network analysis in *Streptococcus pneumonia* and *Salmonella enteritidis* resulted in new biological insights and the selection of genes for further investigation that were not reported in the original experiments. Furthermore, the central roles of GO classes of “regulation of transcription” and “signal transduction by phosphorylation” governed by SP_0798 and cheA in the induction of pathogenesis were shown.

cheA (chemotaxis protein CheA) is a sensor histidine kinase and a member of a two-component system that is primarily involved in “Environmental Information Processing” and “Signal transduction” (KEGG database at http://www.genome.jp/kegg/). According to the Pfam database at http://pfam.sanger.ac.uk/, cheA contains the following domains: PF01584 (CheW-like domain), PF01627 (Hpt domain), PF02518 (Histidine kinase-, DNA gyrase B-, and HSP90-like ATPase), PF02895 (Signal transducing histidine kinase, homodimeric domain), PF09078 (CheY binding), and PF13589 (Histidine kinase-, DNA gyrase B-, and HSP90-like ATPase).

SP_0798 is a DNA-binding response regulator CiaR and a member of a two-component system. According to the Pfam database, SP_0978 contains PF0072 (response regulator receiver domain) and PF00486 (Transcriptional regulatory protein, C terminal). Similar to cheA, SP_0798 is also involved in “Environmental Information Processing”, “Signal transduction” and two-component system (KEGG database). Based on our results and this information, we conclude that SP_0798 and cheA are functionally orthologous in the systems biology context of bacterial pathogenesis.

Based on the above discussion and the similar observed mechanism between *Streptococcus pneumonia* and *Salmonella enteritidis*, we suggest that “Environmental Information Processing” which carries ON by “Signal transduction” and two-component system pathways are conserved pathways used by bacterial pathogens to regulate pathogenicity. In fact, successful pathogens such as *Streptococcus pneumonia* and *Salmonella enteritidis* have developed sensing systems to rapidly recognise external environmental factors and to react more promptly through more efficient signal transduction systems. A two-component system is a head-tail pathway where one member sits outside the cell and the other member is inside the cell and informs the bacteria about environmental signals/changes. Rapid recognition of environmental alterations such as antibiotics and nutrient changes allows bacteria to react more rapidly, increasing the probability of survival. Two-component systems have a confirmed role in bacterial virulence (13,14).

On the other hand, SP_1713 is the major negative regulator in blood infection of *Streptococcus pneumonia*. The fact that SP_1713 has the ability to regulate a large number of other gene ontology terms and dramatically decreases the global transcriptome expression levels in blood, offers a new avenue for the identification of treatments of blood-based infections such as Bacteremia and Sepsis. This example shows how GO network construction can be employed to discover key GO groups and for GO based gene selection.

## Discussion

GO analysis provides a new avenue for a deeper understanding of gene expression and function, which can be exploited in the context of quality-based gene selection strategies (3,4). While other GO web servers (7,15) support gene annotation in model eukaryotes via user submitted gene lists that must match the single source of annotation used by the server, our web server supports all sequenced prokaryotes and viruses and automatically recognizes gene names from all annotation sources.

In contrast to other web servers, our web server provides interactive visual navigation of the hierarchical tree structure of GO groups weighted according to gene expression values at all levels. Furthermore, our server provides dynamic visual reports (using AJAX technology) such as pie charts (to visualize GO group proportions) and bar charts (to compare GO term enrichments versus reference genome based on hyper-geometric distribution), whereas other web servers present this information in text format or rely on visualization capacity provided by other websites including The European Bioinformatics Institute at http://www.ebi.ac.uk/.

The most significant analytical advantage provided by our web server is the ability to compare GO terms across multiple gene samples (or whole genomes) from multiple biological conditions. At present other web servers (7,15) can only compare one sample against a reference genome. Comparative GO analysis is particularly important as a means to identify the underlying biological pathways recruited under different biological conditions. This is essential if one wishes to identify important candidate genes for perturbation experiments.

Unlike other GO web servers that compare one GO term compared to a reference genome at a time (using the Fisher Exact test), our web server can compare all the GO term enrichments from two or more samples (or whole genomes) simultaneously by using robust non-parametric statistical tests. These tests allow the detection of significant changes in GO distribution as a function of experimental conditions.

Finally, our comparative table report takes into account final transcript/protein enrichment to detect GO terms with special enrichment patterns or with specific enrichment fold-change across multiple samples. This helps identify key GO terms and their associated genes based on their expression prevalence. At present, this is a unique analytical approach.

Global transcriptome based GO analysis was achieved by integrating gene expression levels with GO classifications. This allowed us to compare GO enrichment that better reflected the biological reality of the experiments across multiple samples by taking into account the abundance of gene products. This type of comparison was not previously possible, most likely because the predominant eukaryotic GO databases and web servers (7) would not have benefited from such an analysis. Current GO web applications are mostly developed in eukaryotic genomes (5–7) where protein abundance levels are very poorly correlated with gene expression levels, making the need for transcript abundance weighting less useful.

In this report we have presented a method to build an enriched GO regulatory network using public Gene Ontology data. GO regulatory networks from differentially expressed genes can reveal underlying biological pathways (16). In particular the topology of such networks can highlight highly connected/central GO terms and their associated genes, supporting the discovery of candidate genes.

Furthermore, by looking at networks from different bacterial species we can elucidate common biological pathways. Even though we have only implemented GO regulatory networks for bacteria, this type of network could be very effective for eukaryotes as well, particularly for proteomics data. To our knowledge, no current GO web server provides this capability.

We have also demonstrated how to combine a GO regulatory network with gene expression data. The resultant network can be used to study regulatory effects of genes and GOs on each other. For example, by comparing and overlapping multiple GO regulatory networks for the same genes across multiple biological conditions, we can detect areas of the network that confirm or contradict expected regulatory relationships. This can be used as a means to support or question the validity of original transcriptomic data or indicate the existence of unknown environmental effects in experiments. Moreover, by replacing the GO regulatory network’s nodes with their associated genes one can generate a GO-based gene regulatory network (GRN).

Finally, combining GO-based gene regulatory networks with other types of gene regulatory networks (16) (those that are reverse engineered from transcriptome data) such as co-expression networks (17,18) can lead to the discovery of unknown biological entities or biological mechanisms, particularly where such results are contradictory.

Together, the global transcriptomics based GO enrichment and GO regulatory network, developed in this investigation and implemented in the Comparative GO Web application (3,19) can significantly increase our knowledge of bacterial regulatory mechanisms of pathogenesis as well as functional genomic arrangements in newly emerging, highly pathogenic strains.

## Acknowledgments

We would like to greatly thank Dr. Abiodun Ogunniyi, Dr. Layla Mahdi and Prof. James Paton from the Research Centre for Infectious Diseases of The University of Adelaide for their comments and help. We would also like thank Dr. Dan Kortschak for his helpful comments.

## Funding

This work was funded by the University of Adelaide.

## Additional Files

| File Name | File Format | Title of Data | Description of Data |
| --- | --- | --- | --- |
| AdditioanlFile1.xlsx | xlsx | transcriptome | *streptococcus pneumonia* Whole transcriptome Relative Expression Levels Lung vs. Nose (2 Colour Microarray) |
| AdditioanlFile2.xlsx | xlsx | transcriptome | *streptococcus pneumonia* Whole transcriptome Relative Expression Levels Blood vs. Lung (2 Colour Microarray) |
| AdditioanlFile3.xlsx | xlsx | transcriptome | *streptococcus pneumonia* Whole transcriptome Relative Expression Brain vs. Blood (2 Colour Microarray) |
| AdditioanlFile4.xlsx | xlsx | transcriptome | *Salmonella enteritidis* global transcriptome high pathogenic (average of 3 strains) RPKM Counts |
| AdditioanlFile5.xlsx | xlsx | transcriptome | *Salmonella enteritidis* global transcriptome low pathogenic (average of 3 strains) RPKM Counts |
| AdditioanlFile6.xlsx | xlsx | samples | List of Differentially Expressed Genes in Case Study 1 and Study 2 for GO Network Analysis |

